# Cell Tree Rings: the structure of somatic evolution as a human aging timer

**DOI:** 10.1101/2022.12.14.520419

**Authors:** Attila Csordas, Botond Sipos, Terezia Kurucova, Andrea Volfova, Frantisek Zamola, Boris Tichy, Damien G Hicks

## Abstract

Biological age is typically estimated using biomarkers whose states have been observed to correlate with chronological age. A persistent limitation of such aging clocks is that it is difficult to establish how the biomarker states are related to the mechanisms of aging. Somatic mutations could potentially form the basis for a more fundamental aging clock since the mutations are both markers and drivers of aging and have a natural timescale. Cell lineage trees inferred from these mutations reflect the somatic evolutionary process and thus, it has been conjectured, the aging status of the body. Such a timer has been impractical thus far, however, because detection of somatic variants in single cells presents a significant technological challenge. Here we show that somatic mutations detected using single-cell RNA sequencing (scRNA-seq) from thousands of cells can be used to construct a cell lineage tree whose structure correlates with chronological age. De novo single-nucleotide variants (SNVs) are detected in human peripheral blood mononuclear cells using a modified protocol. A default model based on penalized multiple regression of chronological age on 31 metrics characterizing the phylogenetic tree gives a Pearson correlation of 0.81 and a median absolute error of ∼4 years between predicted and chronological age. Testing of the model on a public scRNA-seq dataset yields a Pearson correlation of 0.85. In addition, cell tree age predictions are found to be better predictors of certain clinical biomarkers than age alone, for instance glucose, albumin levels and leukocyte count. The geometry of the cell lineage tree records the structure of somatic evolution in the individual and represents a new modality of aging timer. In addition to providing a numerical estimate of ‘Cell Tree Age’, it unveils a temporal history of the aging process, revealing how clonal structure evolves over life span. Cell Tree Rings complements existing aging clocks and may help reduce the current uncertainty in the assessment of geroprotective trials.

## Introduction

Aging refers to the systematic decline in cellular and organismal function over time. The ubiquity of age-related disease makes chronological age the single most important risk factor for morbidity and mortality [1]. Interventions to slow, delay or even reverse the aging process thus have the potential to mitigate multiple age-related pathologies [2].

To quantify the effectiveness of such interventions it is necessary to have a reliable measure of biological age. Aging timers, or clocks, accomplish this by using specific biomarkers whose states change systematically with chronological age. A variety of biomarker modalities have been studied, particularly epigenetic, but also transcriptomic, proteomic and metabolomic, among others [3, 4]. A necessary step in the development of current aging clocks is to show that the chosen biomarker states are associated with chronological age across a population. This correlation captures the average changes over lifespan and establishes a baseline to which individuals can be compared. A desirable property of these biomarker timers is that they be directly linked to the hallmarks of aging [5]. This potentially allows the biomarker states to be interpreted in terms of the mechanisms of aging.

Genome instability due to somatic mutations is the first hallmark of aging [5]. In blood, mutations can lead to somatic mosaicism and eventually clonal hematopoiesis, where cell populations harbouring particular allele variants outgrow others. Animal models of clonal hematopoiesis have been shown to contribute to disease progression [6]. More generally, diseases characterized by accelerated aging typically involve the increased accumulation of DNA damage [7]. The idea that somatic mutations can drive clonal expansion has stimulated renewed interest in the mutational theory of aging. This represents a new mechanism by which mutations can lead to aging phenotypes [8, 9] and is distinct from earlier proposals which treated absolute mutation burden as a sufficient cause for organismal aging [10]. Given its importance as a driver of aging, it would seem that somatic evolution could form the basis for a new type of aging timer.

Somatic mutations (single-nucleotide variants, SNVs, and copy-number variants, CNVs) are naturally-occurring barcodes [11] that enable phylogenetic inference of cell lineage trees (cell trees from now on). Cell trees are a representation of the mitotic branching order and clonal structure of a sampled cell population [12, 13]. These partial cell trees are subtrees of the whole organismal cell lineage tree which in an adult human consists of tens of trillions of cells [14].

The shape of a tree refers to the ordering and length of its branches and reflects the clonal structure and evolutionary distances between cells.

The central conjecture behind our proposed aging timer is that the structure, or “shape”, of cell trees is a representation of the biological aging process [Csordas, 2019, https://doi.org/10.7287/peerj.preprints.27821v7]. There are two reasons for this hypothesis. The first is that phylogenetic systematics has long shown how genetic distances between species existing today reflect evolutionary changes in the past. It is reasonable then to expect that genetic distances between single cells can be used to infer the somatic evolutionary history of cells, a driver and indicator of aging. The second is that biomedical life history can leave its imprint on the cell tree [15], providing a record of major transitions in the aging process. An additional benefit of cell trees is that they provide an intuitively appealing representation of the dynamics of aging that naturally lends itself to interpretation.

Using human peripheral blood cells from healthy individuals (n=18, age range 21-82 years of age) we have developed a new aging timer called Cell Tree Rings (CTR) with the following characteristics:

1. Naturally occurring somatic single nucleotide variants (SNVs) are used to build cell trees using standard phylogenetic algorithms,
2. SNVs are called directly and de novo from scRNA-seq data from hundreds or thousands of cells,
3. A broad set of tree metrics is used to identify aspects of tree shape that are associated with chronological age using a penalized multiple regression model,
4. The model is used to predict a Cell Tree Age for individuals.

Two different types of phylogenetic algorithms, UPGMA and maximum likelihood, are shown to produce a working Cell Tree Age model, providing extra evidence for the hypothesis. Importantly, the model is validated with public data as an independent test set. The predicted Cell Tree Ages are also shown to correlate with some clinical blood biomarkers, for instance glucose, albumin, leukocytes and monocytes (See *Supplementary Information: Clinical Markers*).

## Methods

### Experimental data and protocol

#### Biological sample collection and isolation of cells

18 blood samples, 5 ml each, have been collected by venipuncture at the Healthy Longevity Clinic (HLC) in Prague, Czech Republic. The samples have been taken with informed consent from healthy patients of the clinic. The Healthy Longevity Clinic Ethical Committee has reviewed and approved the Tree Ring Pilot observational study protocol with the reference number 20220301_001. The age range of the volunteers was 21-82 years old at the time of blood collection, 10 volunteers were males and 8 were females. Samples were processed by the same protocol. In the following the data related to the first 6 samples are detailed. Viable peripheral blood mononuclear cells (PBMCs) were isolated from the collected biological sample. 4 ml of peripheral blood was diluted with 4 ml of 2% Fetal bovine serum (FBS) in Phosphate buffered saline (PBS). Subsequently, 8 ml of diluted peripheral blood was carefully layered on top of 4 ml of a density gradient (such as Lymphoprep^™^) and centrifuged at 300 g for 30 min. The cells were carefully harvested from the interface with a plastic pasteur pipette. Then, another 6 ml of 2% FBS/PBS was added to the cells and centrifuged at 300 g for 8 min discarding the supernatant and resuspending the cells in 1 ml of lysis solution. After one-minute incubation on ice, 4 ml of 2% FBS/PBS was added to the cells and centrifuged at 300g for 5 min discarding the supernatant and resuspending the cells in 1 ml of 2% FBS/PBS. Subsequently, the vitality and concentration of cells was determined through Acridine Orange and Propidium Iodide assay at LUNA Automated Cell Counter. Cell concentration range was between 3.72×10^6^ – 6.35×10^6^ b/ml, and cell viability was between 99.1-99.7%.

#### Labeling the cells with CellPlex

The cells were labelled with molecular tags or CellPlex (according to original protocol CG000391 Cell Labeling with Cell Multiplexing Oligo RevA). Later, a specific volume of each sample was transferred into new 2 ml tubes and, after labeling, the cells were washed 3 times with 2% FBS/PBS (compared to 2 times in the original protocol). After the last wash, the cells were resuspended in 600 μl of 2% FBS/PBS and counted at LUNA. Cell concentration range was between 3.15×10^6^ – 3.95×10^6^ b/ml, and cell viability was between 99.3-99.7%, post labeling.

The samples were pooled proportionally, and the final pool was passed through a 30 μm filter. Finally, the cells were counted and diluted to optimal concentration.

#### Loading and library preparation

Cells were loaded on the Chromium Controller and libraries prepared according to the original protocol CG000390 Chromium Next GEM Single Cell3 v3.1 Cell Surface Protein Cell Multiplexing RevB, aiming for 16000 recovered cells. Some library preparation steps were modified slightly. During cDNA amplification, the polymerization step was extended to 1.5 min. After cDNA purification, the samples were split into two aliquots (A and B) that were processed in parallel and differed only in the implementation of size selection. After fragmentation, double size selection was modified for samples according to Table 1 below.

**Table 1.**
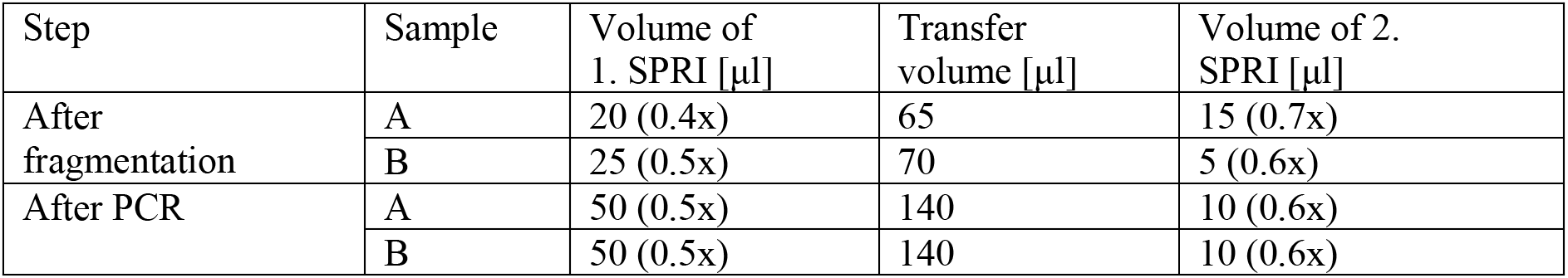
Double-sided size selection using SPRIselect beads.

| Step | Sample | Volume of 1. SPRI [μl] | Transfer volume [μl] | Volume of 2. SPRI [μl] |
| --- | --- | --- | --- | --- |
| After fragmentation | A | 20 (0.4x) | 65 | 15 (0.7x) |
|  | B | 25 (0.5x) | 70 | 5 (0.6x) |
| After PCR | A | 50 (0.5x) | 140 | 10 (0.6x) |
|  | B | 50 (0.5x) | 140 | 10 (0.6x) |

After PCR amplification, both samples were purified using SPRIselect beads according to Table 1. Finally, the quality and quantity of libraries was determined using the Fragment Analyzer and QuantiFluor dsDNA System.

The various chemicals or kits used were Next GEM Chip G Single Cell Kit, Next GEM Single Cell 3’ Gel Beads Kit v3.1, Next GEM Single Cell 3’ GEM Kit v3.1, Dynabeads MyOne Silane, Next GEM Single Cell 3’ Library Kit v3.1, Single Index Kit T Set A, 3’ CellPlex Kit Set A, 3’ Feature Barcode Kit, and Dual Index Kit NN SetA.

#### Sequencing

Library pools were sequenced on an Illumina NovaSeq 6000 using the S4 300- cycle kit and with 150 bp long R2.

### Bioinformatics Processing of the experimental data

The output sequencing files have been processed with Cell Ranger v6.0.2 and the indexed paired-end bam files have been converted into fastq files with bamtofastq v2.30.0. The fastq files and the identified barcode list were used for further processing.

### General schematics of the Cell Tree Rings computational workflow

The Cell Tree Rings computational pipeline involves four consecutive steps, names of the sub-pipelines highlighted in bold.

1. **Tizkit**: Barcode specific calling of SNVs with scSNV and germline filtering.
2. **Tiznit**: Generating fasta files and phylogenetic inference of cell trees.
3. **AgeTreeShape:** Compute tree features and univariate regression on age.
4. **CellTreeAge:** Building multiple penalised regression model and Cell Tree Age prediction. The following four sections provide further details of this workflow.

### Somatic mutation calling de novo from scRNA-seq

We have used scSNV v1.0b [16] to call somatic mutations, specifically single nucleotide variants, directly from 10x Genomics scRNA-seq data through collapsed molecular duplicates to increase mutation coverage. The GRCh38 (hg38) reference human genome build was used for mapping and alignment, specifically GENCODE Release 44, GRCh38.p14. Potential germline variants over 1% of minor allele frequency have been removed using the latest version 110 release of the 1000GENOMES vcf file using the Ensembl ftp directory. The ‘V3’ parameter was set to process 3-prime libraries. The default setting of scSNV has been used with Maximum Variant Allele Fraction set to 0.999. To reduce the number of false positive calls two important consecutive filters were introduced. First, only somatic variants detected by at least 8 different UMIs per barcode have been used, and second, only somatic variants that were present in at least 4 barcodes were considered further for phylogenetic tree generation. The inputs were fastq files and the outputs were sparse SNV count matrices for alternative and reference alleles along with annotated SNV files in csv and vcf format.

### Phylogenetic Tree Inference

The matrices and csv files generated in the previous step are used to generate fasta alignments. Fasta files are generated from all the cells and a subset of the cells with SeqKit v2.1.0 [17]. To track within-sample variability of the trees we generate 5 replicate trees from each sample with each tree being constructed from a random selection of 700 out of the ∼1400 cells from that sample. Because these subsets of 700 cells are partially overlapping with each other, each tree is a pseudo-replicate. We will refer to a pseudo-replicate tree as a “partial tree”

For phylogenetic inference with UPGMA, the R package phangorn v2.8.1 [18] was used with helper functions from the ape package v5.6.2. We used the p-distance (the proportion of sites that differ between a sequence pair) to determine branch lengths by setting the evolutionary substitution model to ‘raw’. The matrix of pairwise distances was computed with the dist.dna function of the ape package. Tree inference provided rooted, ultrametric trees by default. The trees were stored in newick files.

For Maximum Likelihood, IQ-TREE multicore version 2.2.0-beta COVID-edition was used. The substitution model was JC69.

Cell Trees have been visualised with version 1.4.4 of the FigTree tree figure drawing tool.

### Cell Tree Metrics

The tree metrics, or features, can be split into 5 groups based on their technical properties (details in *Supplementary Information: Tree Metrics).* Group I contains spectral tree metrics that are based on the transform matrices of the cell trees which are discrete analogues of the generalized Fourier [19] and Laplacian transforms [20], respectively. Group II contains specialised phylogenetic features focusing on aggregated branch length statistics and their derivatives, including entropy-based metrics. Group III includes well-known general phylogenetic tree statistics used in the biodiversity literature. Group IV focuses on branch length values specifically and generates summary statistics based on the distance matrix between the tips of the tree. Finally, Group V has 2 powergraph based features generating the Laplacian transforms of the square of the tree graphs, similar to Group I.

### Regression Analysis

#### Elastic net regression

To build a predictor model we regress chronological age on 31 tree statistics in addition to Sex, giving a total of 32 features. We allow pairwise interactions between Sex and each tree statistic giving a total of 32+31=63 predictors.

In line with the majority of previous aging timers [21, 22, 23, 3] we employ elastic net regularization [24, 25]. This requires solving

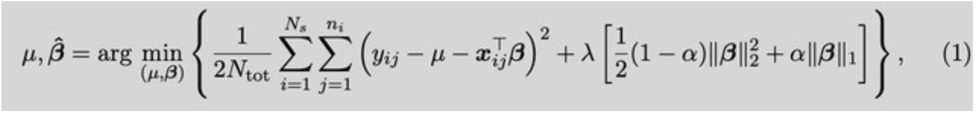

which is a convex program when the hyperparameters λ (the regularization constant) and α (the lasso fraction) are fixed. Here x_ij_ is a predictor and y_ij_ is the chronological age for pseudo-replicate *j* in sample *i*. ***β*** is the vector of regression coefficients, *μ* is the constant offset, *N*_s_ is the number of samples, *n_i_* is the number of replicates in sample *i* and 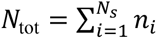 is the total number of replicates across all samples.

Eq. 1 is solved using the elastic-net routine from scikit-learn [26] (version 1.3.0) in Python3 [27] (version 3.11.5).

##### Nested cross validation

To test predictive accuracy using this model we implement a nested cross validation scheme [28, 29]. We used leave-one-out cross-validation in both outer and inner loops: since there are 18 samples this means there were 18 folds in the outer loop and 17 folds in the inner loop (a fold partitions data into a training and test set with each sample assigned to a test set exactly once). All pseudo-replicates from a sample are assigned the same chronological age and are never split across training and test sets.

##### Hyperparameter grid search

Hyperparameters are determined by solving Eq. 1 multiple times, each time with different hyperparameter value combinations. Hyperparameter values are chosen from the sets λ ∈ {0.1,0.3,1,3,10} and α ∈ {0.6,0.7,0.8,0.9,1} in an exhaustive grid search. The hyperparameter combination giving the lowest mean absolute error, as found by cross validation in the inner loop, is chosen as optimal. Once the optimal hyperparameters have been found for a given outer training set, Eq. 1 is solved for one step in the outer loop. The procedure is then repeated for other steps in the outer loop, calculating a new set of hyperparameters each time.

##### Regression coefficients and prediction accuracy

Each step in the outer loop produces a vector of regression coefficient estimates, β̂, and a subset of test sample predictions, *ŷ_i,j_*. Prediction accuracy is calculated from the full set of predicted and chronological age pairs, {*ŷ*)_*ij*_, *y*_*ij*_}.

Performance metrics used are the mean absolute error (mae), median absolute error (mdae), root-mean-squared error (rmse), and Pearson correlation (*r*) defined as follows:

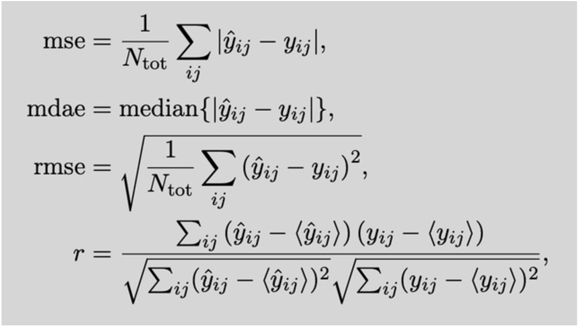

where, for compactness, we write the double sum 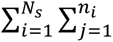 as 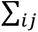

This nested cross validation generates a single age prediction for each pseudo-replicate. Because the outer loop has 18 folds, each regression coefficient is estimated 18 times. Results are shown in Figure 2A, 2B.

##### Testing on public data

A final step in testing the model is to examine its prediction on an external dataset. This involves, in essence, just one step in the outer loop of the nested cross validation where all the 18 HLC samples are used as a single training set, and all the 18 AIDA samples are used as a single test set. Cross validation is still performed in the inner loop to optimize the hyperparameters.

#### Feature pre-selection

The regularized regression procedure described above combines feature selection and coefficient estimation in a single optimization step. This can result in biased predictions since the regularization required to shrink the weaker predictors also shrinks the better predictors [25]. This bias can be mitigated by pre-selecting features in an initial step, prior to elastic net regression [30]. The basic idea is that, by removing some of the weaker predictors prior to regression, the degree of regularization needed in the coefficient estimation step is decreased, thus reducing prediction bias.

Our approach to pre-selecting features is to use the output from the elastic net itself. We take the features selected by the procedure described above and use them in a second (elastic net) regression. This two-step regression approach is similar to adaptive regularization methods where an initial regression step is used to determine feature-specific regularization parameters [31, 32]. In our scheme, the first regression (the selection step) produces a ranking of features based on the magnitude of their regression coefficients. The second regression (the estimation step) is performed with only the top k ranked features. It is the model fit from this second step that is used for prediction.

To determine the optimal value of k, we repeat the estimation step across different k, finding the value that maximizes performance. Technically, the number of features k is a hyperparameter of the model and should be determined in the inner cross-validation loop, along with α and l_1_. This would help account for the uncertainties in post- selection inference [33, 34] and reduce the risk of overfitting. However, for our exploratory purposes, we simply perform the estimation step at several different values of k and use the k that gives the best performance.

We find that feature pre-selection does improve predictability, albeit only slightly (see Supplementary Figure 1 in *Supplementary Information: Tree Metrics*). Thus, for simplicity, our default model uses all features, without pre-selection. Nevertheless, demonstrating that a subset of the features provides as good, if not slightly better, predictability as all the features is helpful for reducing the number of regression coefficients that need to be interpreted (see *Discussion*).

### Processing Public Datasets

The Asian Immune Diversity Atlas (AIDA) public Human Cell Atlas (HCA) scRNA-seq dataset has been used to process candidate samples for model validation [https://data.humancellatlas.org/explore/projects/f0f89c14-7460-4bab-9d42-22228a91f185]. Fastq files have been downloaded from the HCA site. Filtered barcode lists have been directly downloaded from the CellRanger output available at the Chan Zuckerberg CELLxGENE Collections at the following URL: https://cellxgene.cziscience.com/collections/ced320a1-29f3-47c1-a735-513c7084d508 For scSNV, the ‘V2_5P’ parameter was set to process the 10X V2 5-prime libraries.

## Results

### Cell trees

Phylogenetic trees were inferred using the distance matrix algorithm UPGMA and maximum likelihood. To characterise the variability in trees sampled from a single individual in the HLC dataset, we construct trees from 5 subsets of 700 randomly sampled cells rather than a single tree from all ∼1400 cells per individual. Because the sets of 700 cells are partially overlapping, we refer to these “partial” trees as pseudo-replicates. Each partial tree is generated using the same filters as described in *Somatic mutation calling de novo from scRNA-seq* in the Methods section. The regression model fit to the HLC dataset is tested on samples from the AIDA dataset using 5 pseudo-replicate trees, each generated from 700 cells per individual.

For visualisation, Figure 1 shows the complete trees utilising information from all ∼1400 barcoded cells from each individual in the HLC cohort. The circular rendering, which places the root at the centre and the cells around the perimeter, provided inspiration for the name ‘Cell Tree Rings’.

**Figure 1:**
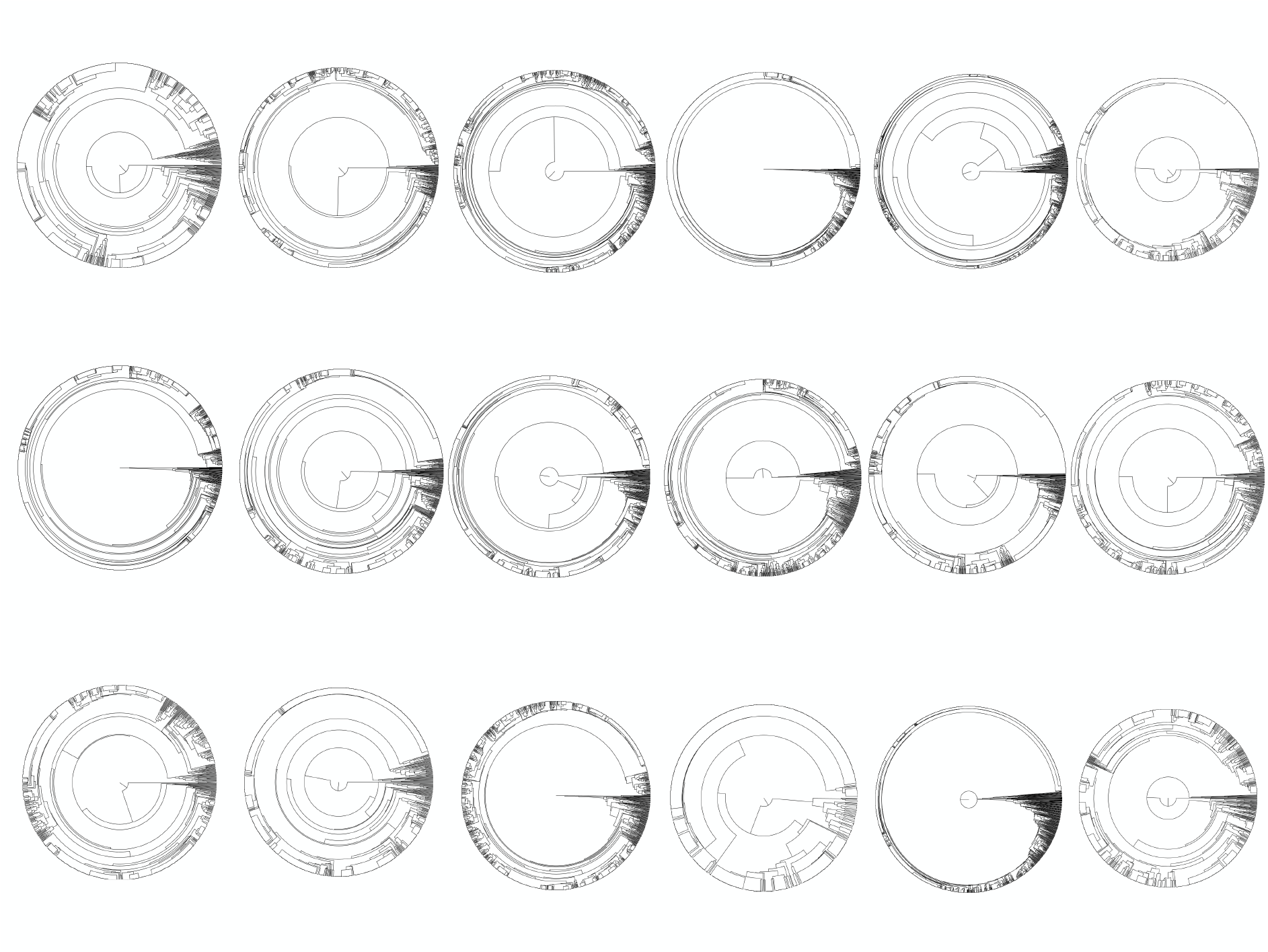
Complete cell trees from each of the 18 HLC participants in the study.

### Cell tree metrics

The central hypothesis of the study is that the shape of cell trees is a measure of biological age. Here shape refers to the combination of topology (branching order) and branch lengths. Topology, in the case of cell lineage trees, corresponds to branching patterns of mitotic division in somatic cells. Branch lengths represent the amount of evolutionary change and are usually defined as the product of mutation rates and a suitable unit of time.

Different tree metrics capture either topology-only, branch-length-only, or a combination of both. We have applied a set of 31 tree metrics to characterize the shape of cell trees built from somatic mutations from human peripheral blood mononuclear cells (See *Supplementary Information: Tree Metrics*). This set of measures comprises both traditional tree metrics used in phylogenetics and some that were developed specifically for this study.

### Model building and Prediction Results

An age predictor has been constructed by regressing chronological age on the 31 tree metrics, along with Sex. Elastic Net regularization and nested leave-one-out cross validation were used to validate and test the model (See *Regression Analysis* in Methods section). The resulting prediction errors, correlation coefficient, p-values and explained variance were estimated by comparing Cell Tree Ages to chronological ages.

Figure 2 shows the performance of the default Cell Tree Age model cross-validated on the 18 samples from the HLC data. In this default model, UPGMA is used for phylogenetic tree inference, 5 pseudo-replicate trees are used per sample, and all 32 features are used in the regression. For this model, *r*=0.813, *p*=0.00004, R^2^=0.660, MAE=7.625, MdAE=4.396, and RMSE=10.760.

**Figure 2:**
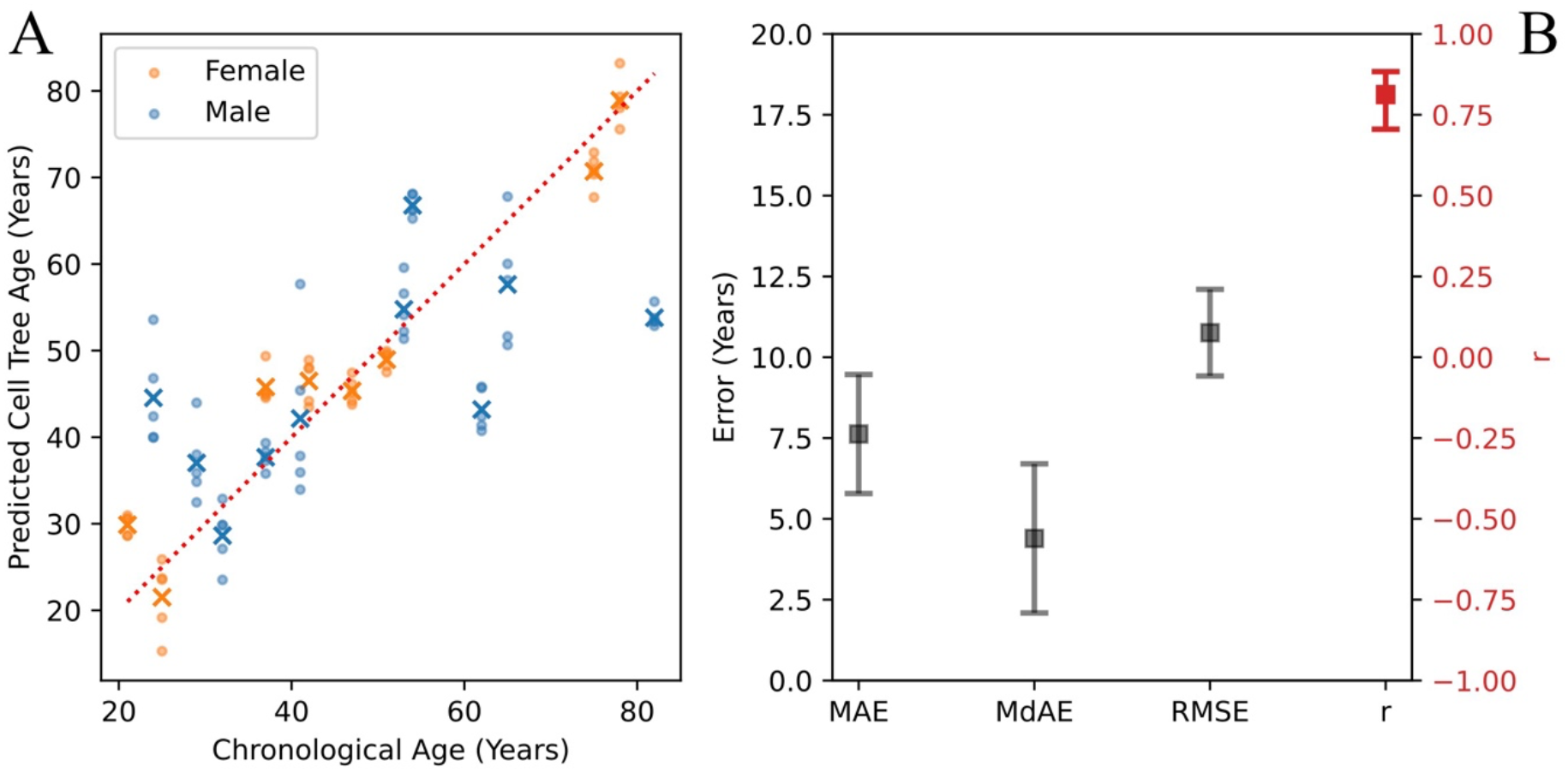
Panel A: Comparison of predicted versus chronological age for 18 human HLC blood samples. Dots represent predictions for each of the 5 pseudo-replicates per sample while crosses represent the mean prediction for each sample. The dotted red line is the reference for perfect prediction (y=x). Panel B: Performance metrics, MAE (mean absolute error), MdAE (median absolute error), RMSE (root mean square error), *r* (Pearson’s Correlation Coefficient).

The Supplementary Information shows how feature pre-selection can slightly improve performance and how using Maximum Likelihood instead of UPGMA for tree inference affects results.

The slope of predicted to chronological age is less than 1, indicating that low ages are systematically overestimated while high ages are systematically underestimated. This is, in large part, due to regularization which biases the regression coefficients towards zero and thus biases predictions towards the mean of the data. Figure 3A illustrates this trend by showing how the age difference (predicted minus chronological age) is positive below and negative above the mean age of ∼47 years. This bias in age prediction can be characterised by a linear trend line (green dashed). It has become customary to refer to the difference between the predicted age and this linear trend line as the “age acceleration” [22], shown in Figure 3B.

**Figure 3:**
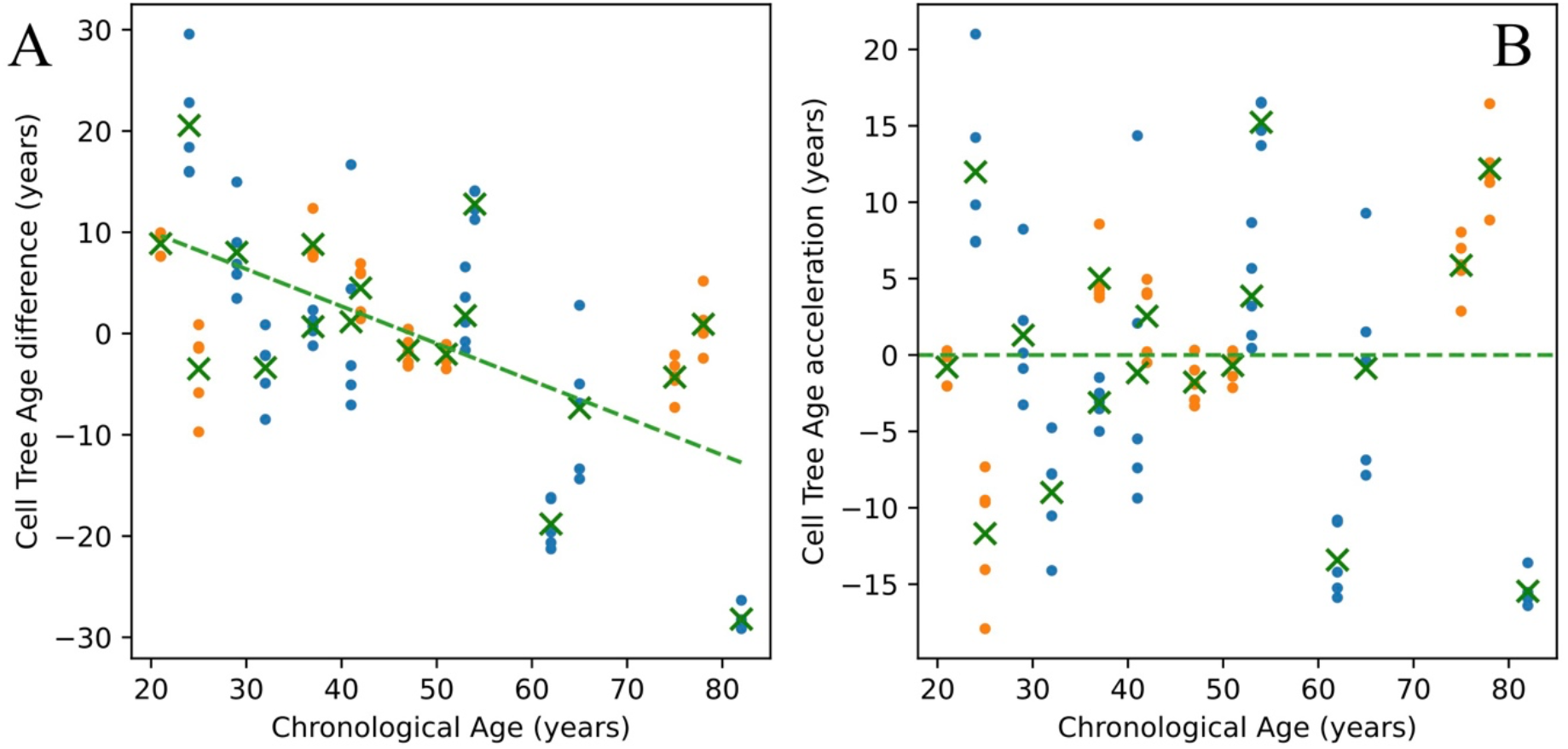
Age difference (Panel A) and age acceleration (Panel B) from the default Cell Tree Age prediction model in Figure 2. The linear trend of predicted to chronological age is given by the green linear. Age difference is predicted age minus chronological age. Age acceleration is the difference between predicted age and the linear fit of predicted to chronological age.

### Public Validation of Cell Tree Age Model

Having developed the Cell Tree Age Model on HLC data, we then tested it on a public scRNA-seq dataset. For this we used 18 peripheral blood samples from the Asian Immune Diversity Atlas (AIDA). This involved 10 females and 8 males ranging in age between 21 – 65 years old. For these data, as with the HLC data, we used UPGMA to generate 5 pseudo-replicate trees from each sample where each pseudo-replicate used 700 cells randomly selected from the sample. The default model was trained on all 18 of the HLC data (see *Testing on Public Data* in the *Regression Analysis* section). The resulting performance metrics were *r*=0.853, *p* = 0.00001, R^2^=0.728, MAE=12.791, MdAE=13.636, RMSE=14.081.

## Discussion

We have shown that cell trees constructed using SNVs from human peripheral blood mononuclear cells can predict chronological age. The SNVs underlying these trees can be directly called from the most accessible single-cell sequencing approach, scRNA-seq. Importantly, the trained Cell Tree Age model was validated on an independent test set and the predicted Cell Tree Age was found to be significantly associated with several blood markers (see *Supplementary Information: Clinical Markers*).

The resulting new molecular aging timer, Cell Tree Rings, requires dozens of cell tree metrics as inputs. Performance of this default model can be improved by ranking the features and regressing on a subset of them (See Feature Pre-Selection in *Regression Analysis* section). Figure 1 of *Supplementary Information Tree Metrics* shows that pre-selection of between 5-12 features improves predictive accuracy. At peak accuracy, using 6 regressors (Sex, ’tipDistNorm’, ’mMax_eigengap’, ’mMaxEigen’, ’AC_2’, ’tipRootPatr’) results in predictions with a median absolute error of ∼3.5 years, a correlation coefficient of 0.9 and ∼82% explained variance. Note that accuracy rapidly declines with fewer predictors such that univariate prediction, even with the single best regressor, has poor accuracy.

Feature selection is useful for identifying the metrics important for prediction. ‘mMax_eigengap’ has been used as a heuristic to identify different modes of divisions or modalities within the tree [20]. ’AC_2’ is the algebraic connectivity of a graph, a well-known feature of graph robustness. Overall, these tree metrics may represent a conceptually new and experimentally verifiable class of quantitative predictors of age.

The Maximum Likelihood method for phylogenetic inference serves as important additional phylogenetic evidence to justify the Cell Tree Age model approach. On the HLC data the best performing model contained 6 predictors with performance metrics r=0.797, p = 0.00007, mae=8.260, mdae=5.654 (See Supplementary Figure 3 *Supplementary Information: Tree Metrics*).

Most importantly, the Cell Tree Age model, trained on the HLC data, when tested on samples from the public AIDA dataset gave an *r* of 0.85 indicating that the model does generalize to existing public data. To further the generalizability, the HLC data came from a Central European cohort, the AIDA data were from an East Asian cohort. However, despite this strong correlation between predicted and chronological age, the clock appears to systematically overestimate age in the AIDA data (see Figure 4). Although feature pre-selection reduces this bias (See Supplementary Figure 2 in *Supplementary Information: Tree Metrics*), there is still a clear overestimate of age.

**Figure 4:**
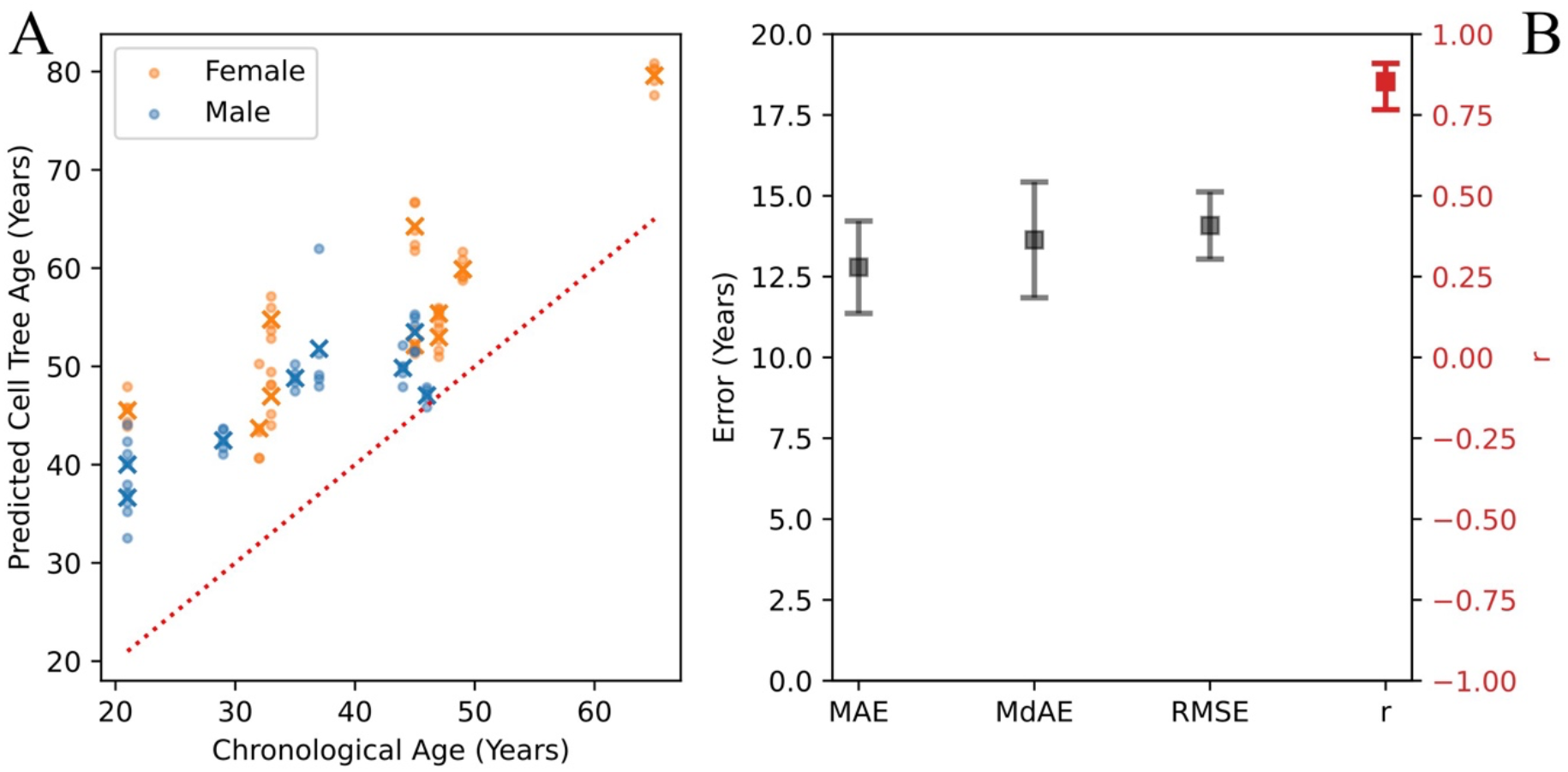
Panel A: Comparison of predicted vs chronological ages for 18 independent human blood samples from the public AIDA dataset. The default model, without feature pre-selection, was trained on 18 samples from the HLC dataset. As in Figure 2, dots represent predictions for each of the 5 pseudo- replicates per sample while crosses represent the mean prediction for each sample. The dotted red line is the reference for perfect prediction (y=x). Panel B: Performance metrics, MAE (mean absolute error), MdAE (median absolute error), RMSE (root mean square error), *r* (Pearson’s Correlation Coefficient).

There are differences between the HLC training set and the AIDA test set that might account for this batch effect. While the HLC data used 10x V3 3’ prime-end libraries, the AIDA data used 10X V2 5’ prime-end libraries. This means that the former method detects variants in the 3’ end region of the genes, including UTR regions, the latter detects variants from the 5’ prime end region, including regulatory elements. Furthermore, the AIDA dataset is primarily restricted to the 20-50 year old age range, with just a single sample over 60. Further investigations are needed to understand this batch effect.

We believe Cell Tree Rings to be the only existing natural barcoding approach that can build larger trees from thousands of cells using somatic mutations called de novo from scRNA-seq data alone as well as from all autosomes, sex chromosomes and the mitochondrial genome combined. Representative cell trees can capture multiple clonal events in different parts of the accessible cellular genomes, reaching a higher resolution in cell population history than possible with targeted genomics approaches alone.

Another advantage of Cell Tree Rings is its potential to extract both lineage histories and gene expression levels from the same cells using a single method and lab protocol. Combining lineage and phenotypic expression information from the same sample is a considerable challenge and existing approaches offer complex solutions combining different protocols [36].

We have also examined the association of Cell Tree Age with 21 clinical blood markers (see *Supplementary Information: Clinical Markers*). Of the 8 markers that show significant associations with Cell Tree or chronological age, 6 of them are better correlated with Cell Tree age than chronological age. These results provide a preliminary indication of how Cell Tree Age could provide valuable clinical indicators. Ultimately, clocks measuring biological age must be better predictors of clinical markers than chronological age. Not all of these associations were related to explicit immune functions, for instance the glucose and albumin associations suggest that if these results are confirmed on bigger cohorts, then Cell Tree Rings might provide a measure of broader multi-tissue and multi-organ age modalities.

### Potential directions for improvement in CTR

The version of CTR reported here represents a basic proof-of-principle. Here we discuss some of the improvements envisioned. The current version of Cell Tree Rings has been achieved with direct and de novo scRNA-seq alone without the aid of any bulk sequencing approach. At a technical level, bulk exome sequencing data can improve true mutation calls at the single cell level and filter out more noise.

In the future, some of this residual variation in predicated age could be explained by individual medical histories or phenotypes, when they become available. In addition, as mentioned above, gene expression information can be extracted from the same cellular barcodes and from the very same genes whose SNVs have been used to generate the cell tree. Efforts are currently underway to see how much of the currently unexplained variation can be accounted for by the cellular phenotypes and the individual medical histories.

Translational geroscience seeks to identify which elements of the aging process are irreversible under current available treatments and which are amenable to modification by existing therapeutic interventions. The tree features involved in Cell Tree Rings may be valuable in diagnosing the influence of a particular intervention by examining its effect on tree shape.

When extending Cell Tree Rings to other tissues an important question is how spatial aspects of the tree can be incorporated. Cellular elements in complex biofluids, such as blood and saliva, have considerable freedom of movement throughout the human body. In contrast, resident cells of compact, solid tissues in the kidney, intestine, liver for instance are under considerable spatial restrictions. The infiltrating immune cells of compact tissues are less constricted spatially than the resident cells, but more constricted than circulating blood cells. It is an open question how tree shape metrics contributing to the age regression model will change in a more restricted spatial environment. Spatial restrictions have been shown to be important in cancer where evolutionary phylodynamic models have been applied to model boundary-driven solid tumor growth [37]. Combining spatial information with the temporal information of cell trees could thus help improve the ability of cell trees to quantify biological age.

### Questions that Cell Tree Rings can help answer

One surprising finding in the phylogenetic tree-aided developmental biology literature is the degree of asymmetry in phylogenetic lineage trees [38, 39]. These studies showed how, at least on the few samples studied, there can be a substantial difference in the number of surviving progeny between offspring of the first or first few cell divisions, the asymmetry reaching sometimes as large as 10:90%. Cell Tree Rings can help quantify this developmental imbalance further and separate it from changes in later life.

Somatic mutations in a small set of (growth and cancer associated) genes have been shown to propagate clones that become dominant in the hematopoietic system of older individuals [35] and have been directly linked to increased risk of cancer and other chronic diseases [40]. Additionally, somatic mosaicism has been shown, in multiple tissues, to rise with age and to predict disease in animal models [6].

From the perspective of designing interventions, it is important to understand which of the somatic mutations are simply passive indicators and which are active drivers of the aging process. In addition, it will be important to establish which can be targeted with clinical interventions.

By providing a way to quantify the different aspects of tree shape, Cell Tree Rings can be used to identify early indicators of clonal hematopoiesis and diagnose why certain individuals display resilience to the effects of somatic mutation and experience reduced chronic age-associated disease.

### Cell Tree Rings and the timescales and convergence of different clocks

When evaluating the effectiveness of different biological aging clocks, it is important to address the question of what the minimal meaningful temporal unit of biological aging is. While clocks with high temporal resolution can evaluate the short-term effects of interventions, these effects can often be difficult to distinguish from physiological noise. On the other hand, lower temporal resolution over longer time windows may miss important short-term signals [41]. Cell Tree Rings captures the long-term dynamics of somatic evolution that relates to the decades-long processes usually associated with aging but is insensitive to processes that are shorter than the characteristic timescale for detecting somatic mutations. It is an open question whether clocks based on different mechanisms and with different time resolutions can be combined and merged, but ultimately the results from different clocks should be reconciled.

Our hope is that Cell Tree Rings may provide a baseline integrative framework for different aging hallmarks and clocks. There are three reasons this may be possible:

First, Cell Tree Rings operate on the fundamental genome-instability level by tracking somatic mutations in hundreds or thousands of single cells. Importantly, it is the use of the tree structure to constrain these mutations that helps to improve their detection accuracy.

Second, Cell Tree Rings is based on a foundational construct, the somatic evolutionary cell tree, that relates the tens of trillions of somatic cells of a human body to each other and to time.

Third, without identifying the damage somatic mutations cause, it is difficult to design healthy longevity therapies and regimens. Cell Tree Rings captures and organizes this basic mutation information at different levels of the tree hierarchy, potentially providing signposts for which interventions are likely to be most effective.

Cell Tree Rings is thus not simply another aging timer. It aims to provide a foundational principle for clocks. There is considerable uncertainty about whether epigenetic aging clocks can inform us about biological age reversals in clinical trials [42]. Adding Cell Tree Rings as a single-cell resolution clock component might mitigate this uncertainty and improve the assessment of geroprotective trials.

## Contributions

A.Cs. conceived the project and wrote the original manuscript. D.G.H. and B.S. edited the manuscript. B.S. produced the core software pipeline. A.Cs., B.S. and D.G.H. designed the study, the methodology and wrote software. A.Cs. and D.G.H. performed the statistical analysis and supervised the study. A.V collected the blood and F.Z supervised the clinical procedure.

T.K and B.T. performed library preparation and scRNA-seq. T.K. wrote the original draft of the Experimental protocol. All authors read and approved the final version of the manuscript.

## Acknowledgements

This work was solely supported by AgeCurve Limited. Special acknowledgement goes to Petr Sramek, of LongevityTech.Fund for providing crucial infrastructure in the Czech Republic. We would also like to acknowledge the patients of Healthy Longevity Clinic, who volunteered to provide blood. Special thanks to Gavin Wilson for insights on the scSNV pipeline.

## References

1. Partridge L, Deelen J, Slagboom PE. Facing up to the global challenges of ageing. Nature. 2018 Sep;561(7721):45-56. doi: 10.1038/s41586-018-0457-8.

2. Kaeberlein M. Translational geroscience: A new paradigm for 21^st^ century medicine. Transl Med Aging. 2017 Oct;1:1–4. doi: 10.1016/j.tma.2017.09.004.

3. Rutledge, J., Oh, H. & Wyss-Coray, T. Measuring biological age using omics data. Nat Rev Genet 23, 715–727 (2022). 10.1038/s41576-022-00511-7.

4. Macdonald-Dunlop E, Taba N, Klarić L, Frkatović A, Walker R, Hayward C, Esko T, Haley C, Fischer K, Wilson JF, Joshi PK. A catalogue of omics biological ageing clocks reveals substantial commonality and associations with disease risk. Aging (Albany NY). 2022 Jan 24;14(2):623–659. Doi: 10.18632/aging.203847.

5. López-Otín C, Blasco MA, Partridge L, Serrano M, Kroemer G. Hallmarks of aging: An expanding universe. Cell. 2023 Jan 19;186(2):243–278. doi: 10.1016/j.cell.2022.11.001.

6. Evans MA, Walsh K. Clonal hematopoiesis, somatic mosaicism, and age-associated disease. Physiol Rev. 2023 Jan 1;103(1):649–716. doi: 10.1152/physrev.00004.2022.

7. Lodato MA, Rodin RE, Bohrson CL, Coulter ME, Barton AR, Kwon M, Sherman MA, Vitzthum CM, Luquette LJ, Yandava CN, Yang P, Chittenden TW, Hatem NE, Ryu SC, Woodworth MB, Park PJ, Walsh CA. Aging and neurodegeneration are associated with increased mutations in single human neurons. Science. 2018 Feb 2;359(6375):555–559. doi: 10.1126/science.aao4426.

8. Vijg J, Dong X. Pathogenic Mechanisms of Somatic Mutation and Genome Mosaicism in Aging. Cell. 2020 Jul 9;182(1):12–23. doi: 10.1016/j.cell.2020.06.024.

9. Massaar S, Sanders MA. The etiology of clonal mosaicism in human aging and disease. Aging and Cancer. 2023 doi: 10.1002/aac2.12061.

10. Szilard L. On the nature of the aging process. Proc Natl Acad Sci U S A. 1959 Jan;45(1):30–45. doi: 10.1073/pnas.45.1.30.

11. Sankaran VG, Weissman JS, Zon LI. Cellular barcoding to decipher clonal dynamics in disease. Science. 2022 Oct 14;378(6616):eabm5874. doi: 10.1126/science.abm5874.

12. Salipante SJ, Horwitz MS. Phylogenetic fate mapping. Proc Natl Acad Sci U S A. 2006 Apr 4;103(14):5448–53. doi: 10.1073/pnas.0601265103.

13. Wasserstrom A, Frumkin D, Adar R, Itzkovitz S, Stern T, Kaplan S, Shefer G, Shur I, Zangi L, Reizel Y, Harmelin A, Dor Y, Dekel N, Reisner Y, Benayahu D, Tzahor E, Segal E, Shapiro E. Estimating cell depth from somatic mutations. PLoS Comput Biol. 2008 May 9;4(4):e1000058. doi: 10.1371/journal.pcbi.1000058.

14. Sender R, Fuchs S, Milo R. Revised Estimates for the Number of Human and Bacteria Cells in the Body. PLoS Biol. 2016 Aug 19;14(8):e1002533. doi: 10.1371/journal.pbio.1002533.

15. Stadler T, Pybus OG, Stumpf MPH. Phylodynamics for cell biologists. Science. 2021 Jan 15;371(6526):eaah6266. doi: 10.1126/science.aah6266.

16. Wilson GW, Derouet M, Darling GE, Yeung JC. scSNV: accurate dscRNA-seq SNV co-expression analysis using duplicate tag collapsing. Genome Biol. 2021 May 7;22(1):144. doi: 10.1186/s13059-021-02364-5.

17. Shen W, Le S, Li Y, Hu F. SeqKit: A Cross-Platform and Ultrafast Toolkit for FASTA/Q File Manipulation. PLoS One. 2016 Oct 5;11(10):e0163962. doi: 10.1371/journal.pone.0163962.

18. 18. Schliep KP. phangorn: phylogenetic analysis in R. Bioinformatics. 2011 Feb 15;27(4):592–3. doi: 10.1093/bioinformatics/btq706.

19. Hicks DG, Speed TP, Yassin M, Russell SM. Maps of variability in cell lineage trees. PLoS Comput Biol. 2019 Feb 12;15(2):e1006745. doi: 10.1371/journal.pcbi.1006745.

20. Lewitus E, Morlon H. Characterizing and Comparing Phylogenies from their Laplacian Spectrum. Syst Biol. 2016 May;65(3):495–507. doi: 10.1093/sysbio/syv116.

21. Hannum, G., Guinney, J., Zhao, L., Zhang, L., Hughes, G., Sadda, S., Klotzle, B., Bibikova, M., Fan, J.-B., Gao, Y., Deconde, R., Chen, M., Rajapakse, I., Friend, S., Ideker, T., and Zhang, K. (2013). Genome-wide methylation profiles reveal quantitative views of human aging rates. Molecular Cell, 49(2):359–367.

22. Horvath, S. (2013). DNA methylation age of human tissues and cell types. Genome Biology, 14(10):3156.

23. Levine, M. E., Lu, A. T., Quach, A., Chen, B. H., Assimes, T. L., Bandinelli, S., Hou, L., Baccarelli, A. A., Stewart, J. D., Li, Y., Whitsel, E. A., Wilson, J. G., Reiner, A. P., Aviv, A., Lohman, K., Liu, Y., Ferrucci, L., and Horvath, S. (2018). An epigenetic biomarker of aging for lifespan and healthspan. Aging, 10(4):573–591.

24. Zou, H. and Hastie, T. (2005). Regularization and variable selection via the elastic net. Journal of the Royal Statistical Society. Series B (Statistical Methodology), 67(2):301–320.

25. Hastie T, Tibshirani R, Wainwright M. Statistical Learning with Sparsity: The Lasso and Generalizations, CRC Press, 2015.

26. Pedregosa, F., Varoquaux, G., Gramfort, A., Michel, V., Thirion, B., Grisel, O., Blondel, M., Prettenhofer, P., Weiss, R., Dubourg, V., Vanderplas, J., Passos, A., Cournapeau, D., Brucher, M., Perrot, M., and Duchesnay, E. (2011). Scikit-learn: Machine learning in Python. Journal of Machine Learning Research, 12:2825–2830.

27. 27. Van Rossum, G. and Drake, F. L. (2009). Python 3 Reference Manual. CreateSpace, Scotts Valley, CA.

28. Varma, S. and Simon, R. (2006). Bias in error estimation when using cross-validation for model selection. BMC Bioinformatics, 7(1):91.

29. Cawley, G. C. and Talbot, N. L. (2010). On over-fitting in model selection and subsequent selection bias in performance evaluation. J. Mach. Learn. Res., 11:2079– 2107.

30. Doherty, T., Dempster, E., Hannon, E., Mill, J., Poulton, R., Corcoran, D., Sugden, K., Williams, B., Caspi, A., Moffitt, T. E., Delany, S. J., and Murphy, T. M. (2023). A comparison of feature selection methodologies and learning algorithms in the development of a dna methylation-based telomere length estimator. BMC Bioinformatics, 24(1):178.

31. Zou, H. (2006). The adaptive lasso and its oracle properties. Journal of the American Statistical Association, 101(476):1418–1429.

32. Zou, H. and Zhang, H. H. (2009). On the adaptive elastic-net with a diverging number of parameters. The Annals of Statistics, 37(4):1733 – 1751.

33. Berk, R., Brown, L., Buja, A., Zhang, K., and Zhao, L. (2013). Valid post-selection inference. The Annals of Statistics, 41(2):802 – 837.

34. Kammer, M., Dunkler, D., Michiels, S., and Heinze, G. (2022). Evaluating methods for lasso selective inference in biomedical research: a comparative simulation study. BMC Medical Research Methodology, 22(1):206.

35. Mitchell E, Spencer Chapman M, Williams N, Dawson KJ, Mende N, Calderbank EF, Jung H, Mitchell T, Coorens THH, Spencer DH, Machado H, Lee-Six H, Davies M, Hayler D, Fabre MA, Mahbubani K, Abascal F, Cagan A, Vassiliou GS, Baxter J, Martincorena I, Stratton MR, Kent DG, Chatterjee K, Parsy KS, Green AR, Nangalia J, Laurenti E, Campbell PJ. Clonal dynamics of haematopoiesis across the human lifespan. Nature. 2022 Jun;606(7913):343–350. doi: 10.1038/s41586-022-04786-y.

36. Lähnemann D, Köster J, Szczurek E, McCarthy DJ, Hicks SC, Robinson MD, Vallejos CA, Campbell KR, Beerenwinkel N, Mahfouz A, Pinello L, Skums P, Stamatakis A, Attolini CS, Aparicio S, Baaijens J, Balvert M, Barbanson B, Cappuccio A, Corleone G, Dutilh BE, Florescu M, Guryev V, Holmer R, Jahn K, Lobo TJ, Keizer EM, Khatri I, Kielbasa SM, Korbel JO, Kozlov AM, Kuo TH, Lelieveldt BPF, Mandoiu II, Marioni JC, Marschall T, Mölder F, Niknejad A, Raczkowski L, Reinders M, Ridder J, Saliba AE, Somarakis A, Stegle O, Theis FJ, Yang H, Zelikovsky A, McHardy AC, Raphael BJ, Shah SP, Schönhuth A. Eleven grand challenges in single-cell data science. Genome Biol. 2020 Feb 7;21(1):31. doi: 10.1186/s13059-020-1926-6.

37. Lewinsohn MA, Bedford T, Müller NF, Feder AF. State-dependent evolutionary models reveal modes of solid tumour growth. Nat Ecol Evol. 2023 Apr;7(4):581–596. doi: 10.1038/s41559-023-02000-4.

38. Bizzotto S, Dou Y, Ganz J, Doan RN, Kwon M, Bohrson CL, Kim SN, Bae T, Abyzov A; NIMH Brain Somatic Mosaicism Network, Park PJ, Walsh CA. Landmarks of human embryonic development inscribed in somatic mutations. Science. 2021 Mar 19;371(6535):1249-1253. doi: 10.1126/science.abe1544.

39. Fasching L, Jang Y, Tomasi S, Schreiner J, Tomasini L, Brady MV, Bae T, Sarangi V, Vasmatzis N, Wang Y, Szekely A, Fernandez TV, Leckman JF, Abyzov A, Vaccarino FM. Early developmental asymmetries in cell lineage trees in living individuals. Science. 2021 Mar 19;371(6535):1245–1248. doi: 10.1126/science.abe0981.

40. Marongiu F, DeGregori J. The sculpting of somatic mutational landscapes by evolutionary forces and their impacts on aging-related disease. Mol Oncol. 2022 Sep;16(18):3238–3258. doi: 10.1002/1878-0261.13275.

41. Gabbutt C, Schenck RO, Weisenberger DJ, Kimberley C, Berner A, Househam J, Lakatos E, Robertson-Tessi M, Martin I, Patel R, Clark SK, Latchford A, Barnes CP, Leedham SJ, Anderson ARA, Graham TA, Shibata D. Fluctuating methylation clocks for cell lineage tracing at high temporal resolution in human tissues. Nat Biotechnol. 2022 May;40(5):720–730. doi: 10.1038/s41587-021-01109-w.

42. Higgins-Chen AT, Thrush KL, Wang Y, Minteer CJ, Kuo PL, Wang M, Niimi P, Sturm G, Lin J, Moore AZ, Bandinelli S, Vinkers CH, Vermetten E, Rutten BPF, Geuze E, Okhuijsen-Pfeifer C, van der Horst MZ, Schreiter S, Gutwinski S, Luykx JJ, Picard M, Ferrucci L, Crimmins EM, Boks MP, Hägg S, Hu-Seliger TT, Levine ME. A computational solution for bolstering reliability of epigenetic clocks: Implications for clinical trials and longitudinal tracking. Nat Aging. 2022 Jul;2(7):644–661. doi: 10.1038/s43587-022-00248-2.

